# SARS-CoV-2 infection dynamics in lungs of African green monkeys

**DOI:** 10.1101/2020.08.20.258087

**Authors:** Emily Speranza, Brandi N. Williamson, Friederike Feldmann, Gail L. Sturdevant, Lizzette Pérez-Pérez, Kimberly Mead-White, Brian J. Smith, Jamie Lovaglio, Craig Martens, Vincent J. Munster, Atsushi Okumura, Carl Shaia, Heinz Feldmann, Sonja M. Best, Emmie de Wit

## Abstract

Detailed knowledge about the dynamics of SARS-CoV-2 infection is important for unraveling the viral and host factors that contribute to COVID-19 pathogenesis. Old-World nonhuman primates recapitulate mild-moderate COVID-19 cases, thereby serving as important pathogenesis models. We compared African green monkeys inoculated with SARS-CoV-2 or inactivated virus to study the dynamics of virus replication throughout the respiratory tract. RNA sequencing of single cells from the lungs and mediastinal lymph nodes allowed a high-resolution analysis of virus replication and host responses over time. Viral replication was mainly localized to the lower respiratory tract, with evidence of replication in the pneumocytes. Macrophages were found to play a role in initiating a pro-inflammatory state in the lungs, while also interacting with infected pneumocytes. Our dataset provides a detailed view of changes in host and virus replication dynamics over the course of mild COVID-19 and serves as a valuable resource to identify therapeutic targets.

## Introduction

A wealth of clinical and laboratory studies have been reported concerning severe acute respiratory syndrome coronavirus 2 (SARS-CoV-2), the causative agent of coronavirus disease 2019 (COVID-19) (Ge et al., 2020; Tay et al., 2020). Among the many unanswered questions that remain, key issues involve the dynamics of SARS-CoV-2 infection, including the identity of the cells that support active virus replication and the immune response to infection. Though multiple cell types in the respiratory tract have been reported to express the critical receptor (ACE2) needed for entry, as well as the protease (TMRPSS2) needed to initiate replication (Qi et al., 2020; Sungnak et al., 2020), it is not clear which of these cell types the virus actively replicates in. One study, using a new algorithm (Viral Track) to detect viral reads in single cell sequencing samples, suggested that both epithelial cells and macrophages contain viral RNA in human bronchoalveolar lavage fluid (BALF) (Bost et al., 2020). However, since the presence of genomic RNA (gRNA) alone does not equate to productive virus infection, this study did not address whether this detection of RNA was the result of virus replication in these cells.

Beyond viral replication dynamics, a thorough study of the host response to infection at the major site of virus replication, the lungs, is needed to better understand potential causes of organ damage and targets for therapeutic intervention. The use of single cell technologies such as multi-parameter flow cytometry and single cell sequencing allow analysis of individual cell states. Sequencing of single cells in BALF (Liao et al., 2020) and upper respiratory tract swabs (Chua et al., 2020) collected from COVID-19 patients detected markers of severe disease in hospitalized individuals. Others have examined the immune response to infection by utilizing single cell sequencing to profile PBMC samples from COVID-19 patients (Wen et al., 2020; Wilk et al., 2020). However, both approaches have limited power to address the dynamics of infection, the host response in the lungs, the main site of virus replication and disease pathogenesis, and neither provides insight into the immune response in the draining lymph nodes.

The limitations of human studies with respect to time of sampling relative to exposure, differences in exposure dose and route, and capacity for deeply analyzing tissues can be overcome using animal models, albeit currently at the cost of not fully replicating the severe disease observed in humans. Indeed, multiple animal models of SARS-CoV-2 infection are being developed to test therapeutics and vaccines and well as to better understand the dynamics of COVID-19 disease progression and the immune response to infection in a time-resolved manner. Non-human primates are commonly used as models for infection and pathogenesis since they often recapitulate human disease. African green monkeys (AGM) are a commonly used non-human primate model for studies of respiratory viruses, including SARS-CoV (McAuliffe et al., 2004). Recently, two studies showed that inoculation of AGM with SARS-CoV-2 results in mild respiratory disease with virus detected in the upper and lower respiratory tract, suggesting this is a suitable nonhuman primate disease model to study viral infection and host response dynamics (Hartman et al., 2020; Woolsey et al., 2020).

To address some of the major questions about SARS-CoV-2 replication and host response, we used traditional virological methods, single cell RNA sequencing, and immunohistopathology. Our combined approach, using inoculation with infectious as well as inactivated SARS-CoV-2 helped determine which cells the virus is replicating in and assess the host response to virus replication. Together, the result is an emerging picture of the viral and immune events associated with mild COVID-19 disease, aiding the understanding some of the dynamics of SARS-CoV-2 infection with very high resolution.

## Results

### sgRNA detection reveals active virus replication in the respiratory tract of SARS-CoV-2 infected AGM

To study SARS-CoV-2 infection and its consequences, two groups of four African green monkeys were inoculated with replication-competent virus, while two control animals were inoculated with virus inactivated by gamma-irradiation. Clinical signs in the control animals were limited to reduced appetite likely as a response to repeated anesthesia (Table 1). Tachypnea was observed in five of eight animals inoculated with infectious SARS-CoV-2. In these animals, disease was mild to moderate and transient, with animals recovering between 5 and 9 days post-inoculation (dpi) (Fig. S1A and Table 1). At 1, 3, 5, 7, and 10 dpi we collected nose, throat and rectal swabs from all animals. Levels of viral genomic RNA (gRNA) in nose and throat swabs were high after inoculation with SARS-CoV-2 and declined over time. Rectal swabs were positive at most time points in one animal with a severely reduced appetite (AGM8) (Fig. 1).

**Table 1.**
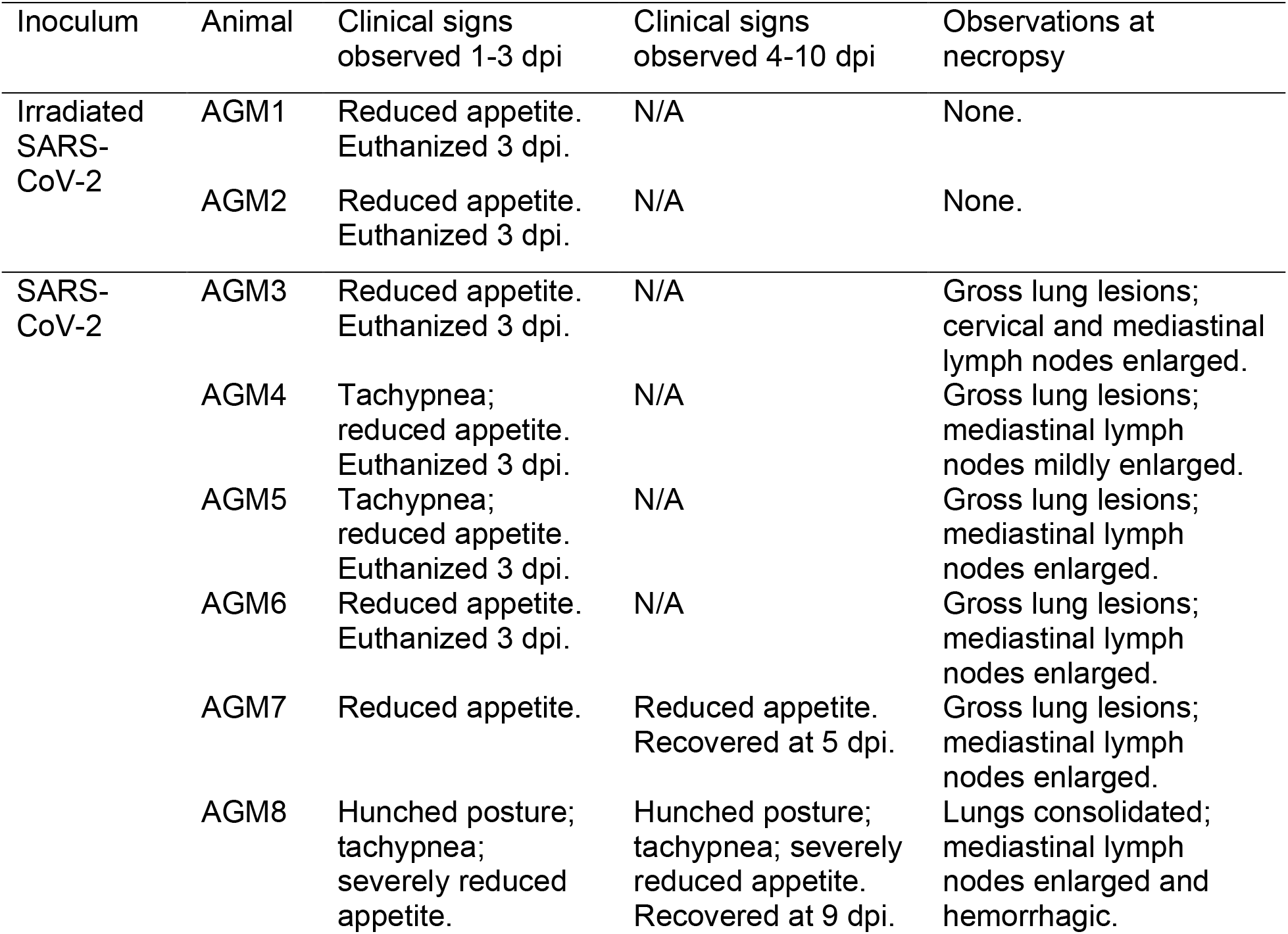
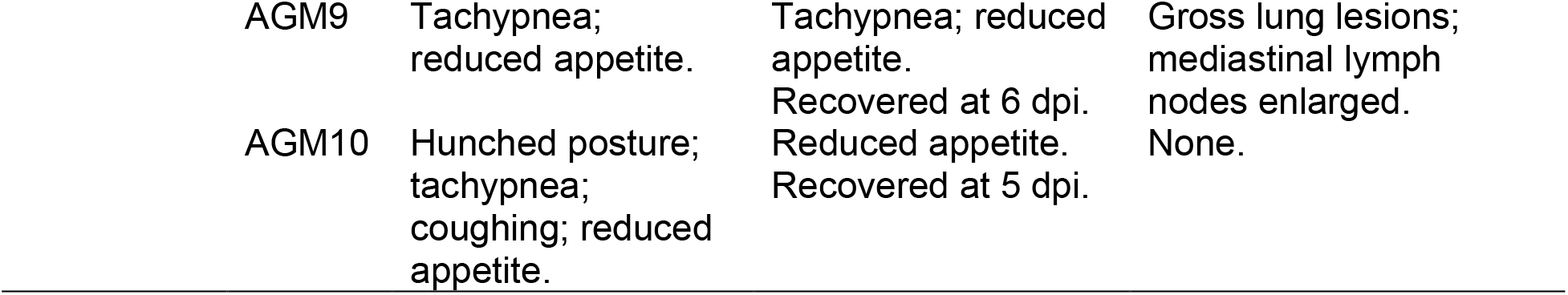
Clinical signs observed in African green monkeys inoculated with irradiated or infectious SARS-CoV-2.

**Figure 1.**
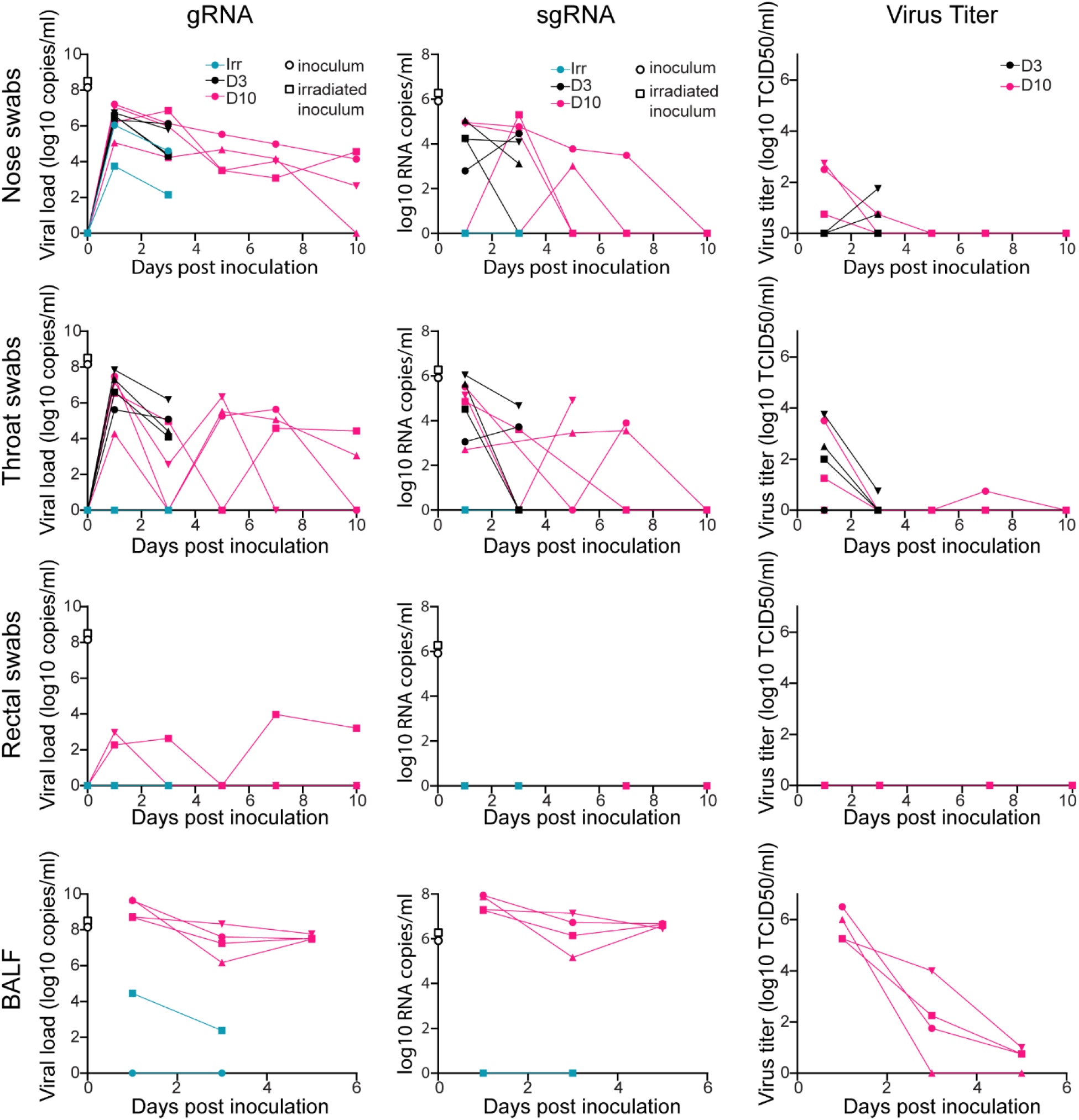
Viral loads and virus titers in swabs and bronchoalveolar lavage fluid. Two African green monkeys (AGM) were inoculated with gamma-irradiated SARS-CoV-2 (n=2). Eight AGM were inoculated with infectious SARS-CoV-2 isolate nCoV-WA1-2020. After inoculation, clinical exams were performed during which nose, throat and rectal swabs were collected; bronchoalveolar lavages (BAL) were performed at 1, 3, and 5 dpi on the four animals remaining in the study through 10 dpi and viral loads and titers were measured. qRT-PCR was performed to detect genomic (left column) and subgenomic RNA (middle column), and *in vitro* virus titration was performed to detect levels of infectious virus (right column) in these samples. Amount of gRNA and sgRNA in the inocula (gamma-irradiated and infectious) is indicated at timepoint zero. Teal: animals inoculated with gamma-irradiated virus; black: animals inoculated with infectious virus and euthanized at 3 dpi; pink: animals inoculated with infectious virus and euthanized at 10 dpi. Where indicated, identical symbols have been used to identify individual animals throughout the figures in this manuscript.

Surprisingly, nasal, although not throat or rectal, swabs from control animals inoculated with gamma-irradiated virus contained high amounts of gRNA at 1 dpi and were still positive at 3 dpi. To determine whether detection of subgenomic RNA (sgRNA) would be able to distinguish between RNA originating from the inoculum from that derived from replicating virus, all swabs positive for gRNA were run in a qRT-PCR to detect subgenomic RNA (sgRNA). Although sgRNA was present at high levels in both inocula, sgRNA could not be detected in swabs collected from control animals but could in nose and throat swabs from animals inoculated with infectious virus (Fig. 1), indicating that sgRNA likely reflects that virus replication occurred. Infectious virus could mainly be detected by virus titration early after inoculation in nose and throat swabs; no infectious virus could be detected in rectal swabs (Fig. 1). As a measure of virus replication in the lower respiratory tract, we collected BALF from the two control animals at 1 and 3 dpi and at 1, 3, and 5 dpi from the four SARS-CoV-2 infected animals euthanized at 10 dpi. gRNA could be detected on 1 and 3 dpi in one of the two control animals; however, sgRNA could not be detected. High levels of gRNA and sgRNA were detected in BALF from the four infected animals, in line with detection of infectious virus through 5 dpi (Fig. 1).

### Assessment of SARS-CoV-2 replication in various tissues

At 3 dpi, the two control animals and four of the SARS-CoV-2 infected animals were euthanized. The remaining four SARS-CoV-2 animals were euthanized at 10 dpi. Upon necropsy, lungs were examined for gross lesions. No abnormalities were detected in the lungs of the two control animals. At 3 dpi, all four animals inoculated with active SARS-CoV-2 showed varying degrees of gross lung lesions and enlarged mediastinal lymph nodes (Table 1 and Fig. S1B). By 10 dpi, one animal did not show gross abnormalities; whereas, the other three animals showed gross lung lesions and enlarged mediastinal lymph nodes (Table 1 and Fig.S1B).

Tissue samples from these animals were assessed for the presence of gRNA and sgRNA. Viral gRNA loads were highest in samples collected from the lung lobes and were higher at 3 dpi than 10 dpi. Despite high levels of sgRNA in lung tissue through 10 dpi, virus could only be isolated at 3 dpi (Fig. S1C and Table S1), indicating that in tissue, sgRNA is a much more sensitive detection method than virus isolation in tissue culture. Analysis of other respiratory tract tissues showed that although gRNA can be detected in all tested sites early after inoculation, sgRNA can be detected consistently only in the trachea and right bronchus (Fig. S1D). Thus, in the respiratory tract of AGM, the lung parenchyma is the main site of virus replication.

We also analyzed tissues of the gastrointestinal (GI) tract for the presence of viral RNA. gRNA alone could be detected in the GI tract of several animals after inoculation with SARS-CoV-2 at 3 dpi and 10 dpi. However, in AGM8, the animal with severely reduced appetite, high levels of both gRNA and sgRNA could be detected in duodenum, jejunum, ileum, cecum and colon (Fig. S1E) and virus was isolated from the ileum and cecum (Table S1). Histologically, the intestinal tract from this animal appeared normal; yet, immunohistochemistry (IHC) revealed epithelial cells containing SARS-CoV-2 antigen in the ileum of AGM8 (Fig. S2A-C).

Histological analysis of the lungs of the two control animals showed no abnormalities (Fig. 2). The lungs of the four animals inoculated with SARS-CoV-2 and euthanized at 3 dpi showed subtle alveolar thickening, indicative of an early inflammatory response (Fig. 2). Viral antigen could be detected by immunohistochemistry in type I pneumocytes and alveolar macrophages of all four animals. The alveolar thickening was still visible in the four animals inoculated with SARS-CoV-2 and euthanized at 10 dpi. Two of these animals showed histopathological changes consistent with interstitial pneumonia frequently centered on terminal bronchioles and early lesions in terminal airways resembling obstructive bronchiolitis (Fig. 2). At this time, viral antigen could only be detected in type I pneumocytes and alveolar macrophages of one of four animals (AGM10). Three of four mediastinal lymph nodes from the 10 dpi samples exhibited a mild to moderate follicular hyperplasia and all four animals’ lymph nodes exhibited rare mononuclear cell immunoreactivity (Fig S2D-E).

**Figure 2.**
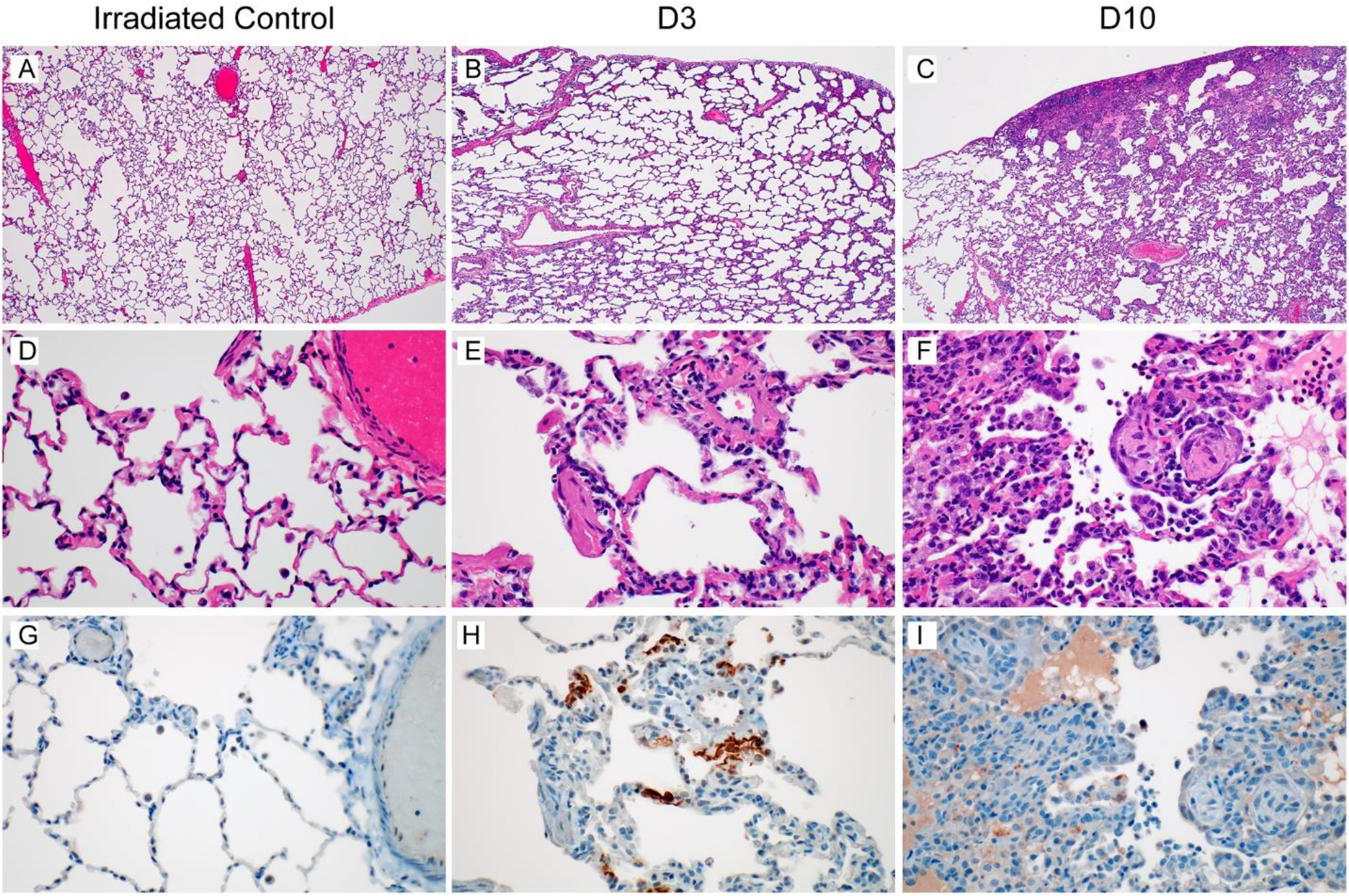
Histological changes in the lungs of African green monkeys inoculated with SARS-CoV-2. African green monkeys were inoculated with gamma-irradiated SARS-CoV-2 (n=2) and euthanized at 3 dpi; eight animals were inoculated with SARS-CoV-2 isolate nCoV-WA1-2020 and four of those were euthanized at 3 dpi, and four at 10 dpi. Histological analysis was performed on lung tissue from all animals. (a) Lungs of animals inoculated with gamma-irradiated SARS-CoV-2 were normal at 3 dpi. (b) Mildly thickened septa were observed at 3 dpi in animals inoculated with infectious SARS-CoV-2. (c) Discrete focus of interstitial pneumonia at the periphery of the lung at 10 dpi. (d) High magnification of normal lung tissue in (a). (e) Alveolar septa are slightly thickened and more cellular at 3 dpi. (f) Alveolar edema (*), type II pneumocyte hyperplasia (arrowhead), increased alveolar macrophages (arrow) and infiltrating lymphocytes and neutrophils are observed at 10 dpi, as well as proliferative nodules associated with terminal airways resembling obstructive bronchiolitis (OB). (g) No SARS-CoV-2 antigen could be detected in lungs from animals inoculated with gamma-irradiated SARS-CoV-2. (h) Cytoplasmic and membrane-associated viral antigen in pneumocytes at 3 dpi. (i) Rare viral antigen could be detected in mononuclear cells, presumably alveolar macrophages, with cytoplasmic debris (arrow) at 10 dpi; background blush is observed in alveolar proteinaceous fluid (*), but pneumocytes do not exhibit immunoreactivity (arrowhead). Magnification a-c 40x, d-i 400x.

### RNA sequencing of single cells from lungs of infected AGM

On the day of necropsy (3 dpi for the two control animals inoculated with gamma-irradiated virus, 3 and 10 dpi for the eight animals that received SARS-CoV-2), we collected sections of the lungs of each animal that contained an active lesion, except in animals where gross lung lesions were not observed at necropsy (Table 1). These sections were processed directly following necropsy into single cell suspensions and single cell RNA sequencing was conducted using cDNA generated immediately without freezing or fixation of the cells, thereby allowing collection of whole cell data. This allowed for high-quality single cell data to be collected with a high fraction of reads in cells (> 80%) and a low fraction of cells enriched in mitochondrial genes (< 5%). Uniform manifold approximation and projection (UMAP) was used to display the data with each cell annotated with its likely cell identity (Fig. 3A). For the latter analysis we developed a correlation-based method that uses transcriptional profiles from annotated lung tissue and determines the most likely identity for an individual cell or a cluster of cells (see methods). To validate the results, we built a marker gene set for each cell type of interest, showing that the gene expression profile matched the cell type annotation (Fig. 3B). Using these assignments, we compared the percentage of cells in each sample belonging to each annotation and found a trend towards an increase in plasma cells as well as an increase in the number of pneumocytes and dividing cells as infection progressed (Fig. S3). This is consistent with the histology results showing an influx of inflammatory cells and type II pneumocyte hyperplasia at 10 dpi (Fig. 2F).

**Figure 3.**
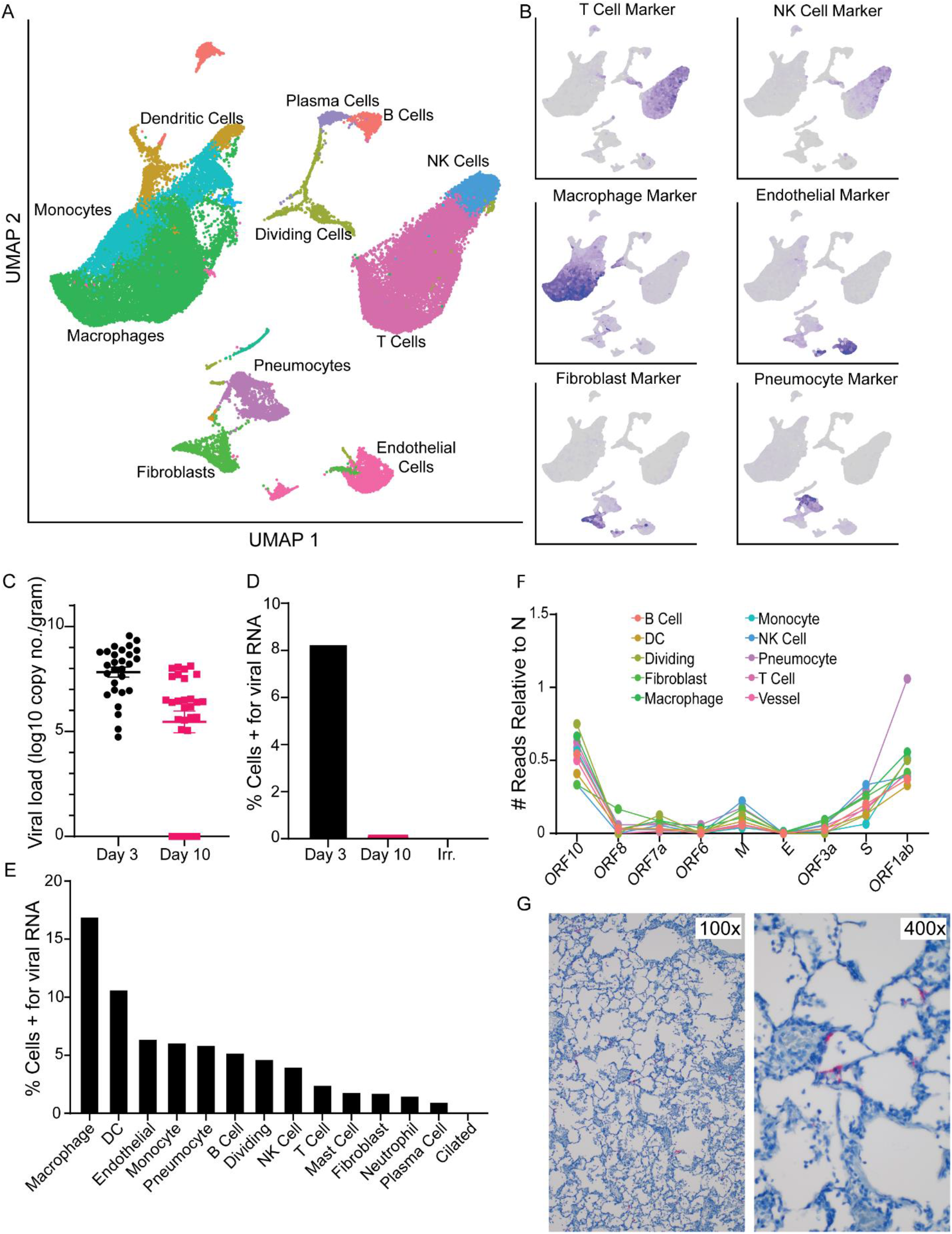
Single cell sequencing and viral dynamics in lung tissue. A. UMAP projection of single-cell RNA sequencing data from whole lung sections from all 10 animals combined. Each point is an individual cell; colors are based on cell type annotation. Cell names are shown next to their largest cluster. B. Validation of cell type identities using marker gene sets. The darker the purple in each cell the higher its expression of the marker set. Grey means the cell did not express any genes in the marker set. C. Viral load information from the lungs via qRT-PCR for gRNA, grouped for all lobes across all animals. D. Percentage of cells identified by single cell RNA-sequencing that are positive for any reads aligning to the viral genome by days post inoculation. E. Percentage of cells from the 3 dpi samples positive for any reads aligning to the viral genome grouped by cell type. F. The number of cells grouped by cell type (colored to match the UMAP in A) with reads aligning to various other locations across the viral genome, all normalized to the number of cells expressing N. Genes are ordered from the 3’ to 5’ end of the SARS-CoV-2 genome. G. ISH for viral S RNA in lung tissues at 3 dpi (viral RNA stains red) at 100x magnification and 400x magnification.

Since SARS-CoV-2 is a poly-adenylated virus, with such modification of both the genome (gRNA) and the transcripts (sgRNA), we could map reads to both the AGM and the SARS-CoV-2 genome and determine which cell types contained viral RNA. The scRNA-Seq data showed a similar pattern to the qRT-PCR data described with the highest percentage of positive cells at 3 dpi, and a decrease by 10 dpi (Fig S1C); no viral RNA was detected in the animals inoculated with gamma-irradiated virus (Figure 3C and D). We then parsed these data down to individual cell types. At 3 dpi, viral RNA could be detected in several cell types, with the macrophage population having the highest percentage of cells positive for viral RNA (Figure 3E). The viral RNA in these cells could be due to virus replication, phagocytosis of infected cells, abortive infections, or having virus particles partitioned with cells during GEM creation in the 10X genomics processing steps. To discriminate between cells supporting active virus replication vs. those containing viral RNA due to other processes, we looked at the distribution of reads across the genome. Due to the 3’ bias of sequencing with the 10X platform, we saw an expected enrichment of reads in the 3’ end (Figure S4A), with a bias to the location of the N gene in the genome. Additionally, in the pseudo-bulk data read pileups, we were able to detect small enrichments of reads in specific areas at the most 3’ end of all the transcripts, including ORF1ab, as well as pileups in the first 5000 base pairs (bp) of the ORF1ab region (Figure S4B). These points of enrichment in the first 5000 bp matched to locations of non-canonical sgRNA formed by a jump of the RNA-dependent RNA polymerase in ORF1ab to N in the SARS-CoV-2 transcriptome and are suggestive of actively replicating virus (Kim et al., 2020). Based on these findings, we examined each cell type for reads showing such evidence of subgenomic transcripts involving a region other than N. Only one annotated cell type, pneumocytes, had a similar proportion of cells that were positive for the N gene and for ORF1ab that was also an abnormally high ratio of ORF1ab to N compared to all other cell types (Fig. 3F). This suggests that despite many cell types containing viral RNA, the pneumocytes were likely the dominant cell type supporting productive viral replication. To examine this hypothesis further, we performed IHC and in-situ hybridization (ISH) on lung tissues at 3 dpi. Although along with pneumocytes, a few macrophages were positive for viral antigen by IHC (Fig. 2H), only pneumocytes were positive by ISH against the viral genome, consistent with the notion that this is the only cell type analyzed supporting active virus replication (Fig. 3G).

### Infection-related changes in lung cell transcriptional states

To gain insight into the biological effects of SARS-CoV-2 infection on diverse cells in the lungs, we examined the transcriptional signatures in cells recovered from animals under each of the viral-exposure conditions (gamma-irradiated 3 dpi, SARS-CoV-2 3 dpi, and SARS-CoV-2 10 dpi). To this end, we developed a method in which each population of cells is examined for a clustering bias in a pair-wise comparison along the various significant principal components. We then used the principal component information to determine gene signatures driving the differences. Using this method, we found that macrophages showed large clustering biases in all the comparisons suggesting they had the biggest transcriptional shift over the course of infection (Fig. S5). The 10 dpi samples, likely due to increases in cell numbers as described above, also showed changes in plasma cells and pneumocytes (Fig. S3 and Fig. S5).

The macrophage analysis was extended by selecting and re-analyzing cells that had either a macrophage or monocyte-like phenotype and further classified to cells with a more tissue-resident phenotype (i.e. alveolar macrophages) or more monocyte-derived phenotype (interstitial macrophages and monocytes) using the expression of MARCO gene (Figure 4A). A comparison of the percentage of the MARCO+/MARCO-cells showed that at 3 dpi there was a large influx of monocyte-derived cells as most of the macrophages detected at that time are MARCO-. This shift began to normalize by 10 dpi as the animals recovered (Fig. 4B).

**Figure 4.**
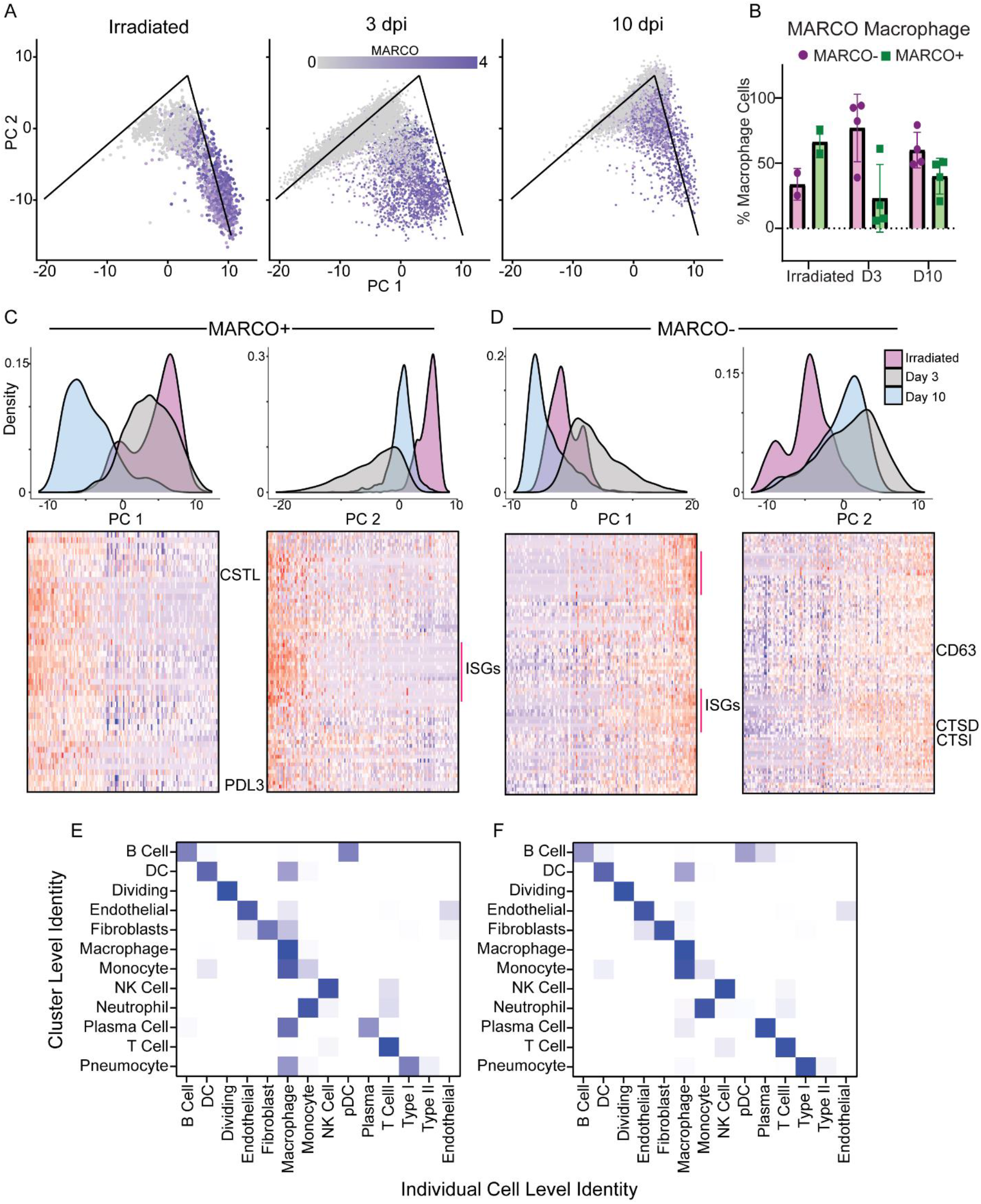
Changes in macrophage populations in the lungs during SARS-CoV-2 infection. A. Principal component analysis of the macrophages. The x-axis is pc1 and the y-axis is pc2. The graph is split by the experimental groups. Each point is an individual cell and is colored based on the expression of MARCO with grey representing zero expression and the dark purple represent increased expression. The lines on the PC graphs are for reference across the samples and represent matching locations. B. The percentage of macrophages that are MARCO + (green) or MARCO – (purple) across the three different conditions (x-axis) C. The MARCO+ macrophages principal component analysis (density plots) grouped by the different conditions. Blue: animals inoculated with gamma-irradiated virus and euthanized on 3 dpi; grey: animals inoculated with SARS-CoV-2 and euthanized at 3 dpi; pink: animals inoculated with SARS-CoV-2 and euthanized at 10 dpi. The left density plot is pc1 with the heatmap below showing the individual cells (columns) sorted based on their location along pc1 with top genes showing high correlation along that principal component clustered (rows). Darker blue means low to no expression and a dark red color means high expression. A similar graph, but for PC2 is on the right. A few of the gene names are noted just to the right of the heatmaps. D. A similar set of graphs as in panel C but for the MARCO-cells. E. Showing the comparison between the individual cell identity (columns) and the cluster level identity (rows) based on the unbiased identification algorithm at 3 dpi for SARS-CoV-2-infected animals. The colors represent the % of individual cells in the cluster that match the identified phenotype. White means no individual cell in that cluster match the phenotype; whereas the darker the blue the higher the percentage of matching cells. F. A similar graph to E at 10 dpi for SARS-CoV-2-infected animals.

To identify the major pathways whose genes are responsible for the analytical differences between these two cell populations, we performed a similar sub-cluster analysis on the MARCO+/MARCO-cells. For the MARCO+ cells, there was a transcriptional shift at 10 dpi along principal component 1 (PC1) (Fig. 4C). Pro-phagocytic gene sets were enriched along this component, suggesting these macrophages were still active in clearing cell debris from the infected lungs potentially explaining why the macrophages stain for viral antigen at this timepoint (Fig. 2I). Comparatively, along PC2, we see a shift mostly at 3 dpi, and only a slight shift at 10 dpi. The genes associated with this component are enriched for pro-inflammatory genes such as interferon-stimulated genes (ISGs) (Fig. 4C). In distinction to the MARCO+ cells, for MARCO-cells, along PC1 we saw a shift in the samples at 3 dpi, mostly with pro-inflammatory genes (Fig. 4D). Along PC2 in the MARCO-cells, we see again the pro-phagocytosis genes associated with the endosomes, as well as migratory genes (CD63). This was observed both at 3 and 10 dpi (Fig. 4D), suggesting that at 3 dpi, the macrophages are in an inflammatory state that is beginning to resolve by 10 dpi, and that the MARCO+ macrophages do not increase their expression of pro-phagocytic genes until later in infection.

To assess interactions between macrophages and other cell types, we looked at a combination of the cluster level identity as compared to the individual cell level identity. Using the same cell identity algorithm, and the clusters calculated by Seurat (Stuart et al., 2019) we looked within individual clusters to identify which cells had a strong macrophage gene marker. At 3 dpi, the clusters associated with pneumocytes, fibroblasts, and endothelial cells all had strong gene markers for macrophages, suggesting these cells were being actively phagocytosed by the macrophages (Fig. 4E). This signature was mostly absent by 10 dpi, possibly because the MARCO+ macrophages are picking up more dead cells and cellular debris by this time rather than whole cells with replicating virus (Fig. 4F).

### Mediastinal lymph nodes are in an inflammatory state at 3 dpi

To relate these virus replication and inflammatory changes in the lungs to changes in secondary lymphoid tissues, we sampled cells from the mediastinal lymph nodes. We collected whole lymph nodes from animals at the time of necropsy and prepped the samples along with the lung tissue. In the lymph nodes, we were not able to detect any viral RNA by single-cell RNA sequencing, despite a few mononuclear cells staining for viral antigen by IHC (Fig. S2E). This could be due to low abundance of RNA, low sensitivity of the scRNA-Seq assay in its ability to detect lowly abundant transcripts, or an inability to capture high abundance of non-lymphocyte populations from the lymph node when generating cell suspensions (Gerner et al., 2012). Using an annotation strategy like that used for lung cells, we could detect most of the major cell populations of the lymph node, with most cells annotating as T or B lymphocytes (Fig. 5A). Again, as with the lung, the annotated cell types expressed known marker genes associated with their phenotype (Fig. 5B). The largest transcriptional change observed across all the cell types was an increase in interferon responsive genes at 3 dpi suggesting an inflammatory state in the draining lymph nodes. This transcriptional profile was resolved by 10 dpi and was absent in the lymph nodes from animals that received gamma-irradiated virus (Fig. 5C).

**Figure 5.**
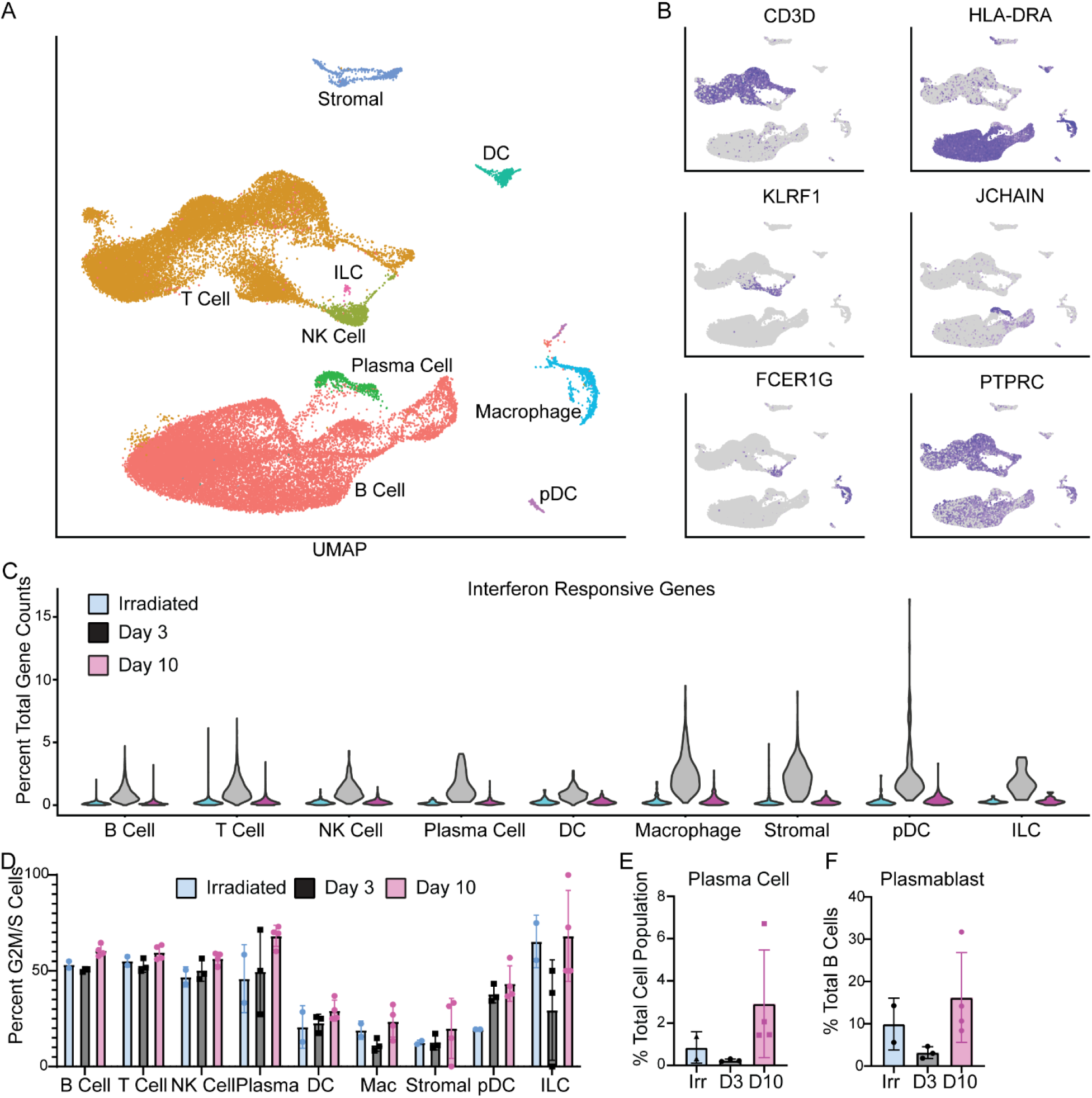
Single cell sequencing of mediastinal lymph nodes. A. UMAP projection of single cell sequencing data from lymph nodes of all 10 animals combined. Each point represents an individual cell and cells are colored based on their cell type. The names of the cell types are placed next to their largest cluster. B. Single gene expression to validate cell type identifications in A. For each graph in this panel, grey means the cell had zero expression and the darker the purple, the higher the expression of that gene. C. percent of total gene counts for each cell for a subset of interferon responsive genes (y-axis). The x-axis is split by the cell types and further divided by sample: blue represents animals inoculated with gamma-irradiated virus and euthanized at 3 dpi; grey represents animals inoculated with SARS-CoV-2 and euthanized at 3 dpi; pink represents animals inoculated with SARS-CoV-2 and euthanized at 10 dpi. Individual cells are removed for clarity and only densities of cells across the populations are shown in violin plots. D. Percentage of each cell population (x-axis) that is actively dividing (stage G2M or S) as determined by a profile of gene expression. Each point is an individual animal and bars represent the mean and standard deviation of the samples. E. Percentage of plasma cells in each animal’s sample compared it’s total cell numbers. F. Percentage of plasmablast cells in each animal’s sample compared to the total B cell population.

There was an increase in the percentage of dividing cells (cells identified as in G2M or S-phase) among B-cells, NK-cells, plasma cells, and pDCs as infection progressed (Fig. 5D). This is consistent with histology findings of lymph node hyperplasia later at 10 dpi (Fig. S2D). Additional changes in cell populations included an increase in plasma cells and plasmablasts by 10 dpi (Fig. 5E). Interestingly, the latter was also seen in the animals that received the gamma-irradiated virus at 3 dpi (Fig. 5F) suggesting the development of an adaptive immune response to the virus. The development of plasmablasts at 3 dpi in animals that received SARS-CoV-2 is lacking, potentially due to the inflammatory state of the lymph node. When looking at which cell type showed the strongest transcriptional changes, we found that even at 10 dpi, macrophages showed the strongest transcriptional shift when compared to samples from animals inoculated with gamma-irradiated virus (Fig. S6A). Within the macrophage population, we saw an upregulation in a small subset of macrophages with markers for fully mature monocyte-derived macrophages such as CHIT1 at 10 dpi (Fig. S6B-C).

## Discussion

This study used large scale single cell sequencing in a non-human primate model of SARS-CoV-2 infection. The benefit of using animal models to study SARS-CoV-2 lies in the ability to collect time-resolved datasets of the lungs instead of being limited to sampling at terminal timepoints (Menter et al., 2020; Tian et al., 2020). Using traditional virological methods as well as single-cell sequencing we have begun to parse out the infection dynamics which occur as the disease progresses and virus is eventually cleared in the African green monkey model of mild COVID-19.

One informative aspect of this study was the inclusion of two animals inoculated with gamma-irradiated SARS-CoV-2 which renders the virus unable to replicate (Feldmann et al., 2019). This enabled us to directly compare the detection of viral genomic RNA from replicating vs. non-replicating virus. We found that sgRNA, though present at high levels in the inoculum, was not present in the swab samples collected at 1 and 3 dpi despite evidence of gRNA from the gamma-irradiated virus, indicating that sgRNA is less stable than gRNA and will degrade quickly without replicating virus generating more such RNA. Such viral dynamics could be due to gRNA being protected and encapsulated or secondary structures which help prevent degradation (Wakida et al., 2020). This has potential implications in patient testing as positive gRNA results by PCR may not represent replication-competent SARS-CoV-2, especially since the highest amount of gRNA detected in the samples from the animals inoculated with irradiated virus were found in the nasal swabs currently used for most patient diagnostics. Our data suggests that PCR-based tests which specifically target SARS-CoV-2 sgRNA may provide a more realistic signature of replicating virus than detection of gRNA, improving models to determine patient infectivity (Bullard et al., 2020; Huang et al., 2020). Since animals cleared sgRNA faster than gRNA the duration of time PCR-positive patients needs to remain in isolation could potentially be reduced.

Using the AGM model, we have been able to explore the dynamics of SARS-CoV-2 infection at the time of peak disease in the lungs where most virus replication is occurring. Due to the nature of the 10X genomics platform and the poly-adenylation of the SARS-CoV-2 genome and sgRNA, we were able to determine which cell types were positive for viral RNA in a method similar to the one used by (Bost et al., 2020), though we did not include any other alignments beyond SARS-CoV-2. Importantly, we were able to detect read enrichment across the whole genome of the virus, though the strongest enrichment was at the most 3’ end of the genome, followed by the 5’ leader TRS. This is consistent with the presence of both gRNA and sgRNA in the lungs at 3 dpi. Further investigation into the distribution of reads across the genome suggests that productive virus replication is mostly occurring in infected pneumocytes, though the macrophage population contained the highest percentage of cells positive for viral RNA. A potential reason that the macrophages could have such high viral RNA and immunoreactivity is through phagocytosis of virus particles or infected cells or through abortive replication. Interestingly, abortive replication could result in the production of aberrant replication products that induce a proinflammatory response as has been shown for influenza A virus in the lungs (Brandes et al., 2013; Te Velthuis et al., 2018).

In the conditions examined here, macrophages appeared to be the major drivers of inflammation in the lungs. This is especially interesting as it has been suggested through the study of healthy human tissue that macrophages interact with ACE2 expressing cells at higher frequencies than other populations (Qi et al., 2020). Both the resident and monocyte-like macrophages at 3 dpi were enriched for pro-inflammatory genes likely caused by having higher titers of virus in the lungs driving the inflammatory state. Interestingly, the MARCO+, or tissue-resident, and MARCO-, or monocyte-derived, macrophage populations showed distinct patterns of gene expression in the 3 and 10 dpi samples. The MARCO+ cells were enriched for ISGs at 3 dpi, but enriched in pathways associated with the endosomes/lysosomes which are up-regulated during a pro-phagocytic period (A-Gonzalez et al., 2017) only at 10 dpi. Only the MARCO-cells at 3 dpi were enriched in genes associated with migration and endosomes/lysosomes, suggesting that, at 3 dpi, macrophages have very different responses to infection based on the cell origin (tissue-resident or recruited) and are likely performing different roles during infection. Further investigation into which macrophage populations are responsible for innate immune host defense and for clearing up the lung environment of cellular and other debris at different stages post-inoculation may help lead to unravel the causes of lung cell damage and indicate potential targets for therapeutic intervention.

Interferon stimulated genes (ISGs) represented dominant responses in monocyte/macrophage populations in both the lung and lymph nodes in the early stages of infection but are reduced by the time infection is cleared. In contrast to the AGM model, the role of the IFN response has been proposed as a driver of disease if it is induced with delayed kinetics relative to peak virus replication (Channappanavar et al., 2016; Channappanavar et al., 2019). Single-cell approaches in human PBMCs and BAL from patients stratified by COVID-19 severity have revealed that type I, II and III IFN is sustained at higher levels in severe patients, while IFNs declined in moderate patients over time from symptom onset (Liao et al., 2020; Lucas et al., 2020; Wilk et al., 2020). In the AGM model, monocyte recruitment and ISG responses were temporally controlled along with virus replication and were diminished in the lungs during recovery and not sustained in the lymph nodes beyond the stage of peak virus replication. Therefore, this model reflects effective viral control and reveals transcriptional signatures within tissues associated with protective responses.

Together, the data reported here provide unique insights into the dynamics of SARS-CoV-2 infection over time in the lungs and associated secondary lymphoid tissue, the identity of the cells hosting replicating virus, and the transcriptional changes in these and other cells during this infectious process. Although the AGM model did not result in severe disease upon inoculation with SARS-CoV-2, it did allow for a deeper understanding of virus replication and host response dynamics along with gene signatures associated with a successful resolution of infection which can be used to inform development of host-targeted therapeutics.

## Acknowledgements

The authors would like to thank Greg Saturday, Dana Scott, Chad Clancy and all RMVB animal caretakers for help with the animal experiment, Anita Mora for help with figure preparations and Ronald N. Germain for helpful discussions. This work is supported by the Intramural Research Program of NIAID, NIH.

## Author contributions

Conceptualization, E.S., V.J.M., H.F., S.M.B and E.d.W.; investigation E.S., B.N.W., J.L., F.F., G.S., L.P.-P., K.M.-W., B.J.S., C.M., A.O., C.S. and E.d.W.; writing-original draft, E.S. and E.d.W.; writing, review and editing, all authors.

## Declaration of interests

The authors have no competing interests to declare.

## Methods

### Ethics and biosafety statement

All animal experiments were approved by the Institutional Animal Care and Use Committee of Rocky Mountain Laboratories, NIH and carried out by certified staff in an Association for Assessment and Accreditation of Laboratory Animal Care (AAALAC) International accredited facility, according to the institution’s guidelines for animal use, following the guidelines and basic principles in the NIH Guide for the Care and Use of Laboratory Animals, the Animal Welfare Act, United States Department of Agriculture and the United States Public Health Service Policy on Humane Care and Use of Laboratory Animals. African green monkeys were housed in adjacent individual primate cages allowing social interactions, in a climate-controlled room with a fixed light-dark cycle (12-hr light/12-hr dark). Animals were monitored at least twice daily throughout the experiment. Commercial monkey chow, treats, and fruit were provided twice daily by trained personnel. Water was available *ad libitum*. Environmental enrichment consisted of a variety of human interaction, manipulanda, commercial toys, videos, and music. The Institutional Biosafety Committee (IBC) approved work with infectious SARS-CoV-2 strains under BSL3 conditions. Sample inactivation was performed according to IBC-approved standard operating procedures for removal of specimens from high containment.

### Study design

To evaluate the pathogenesis of SARS-CoV-2 in African green monkeys, eight adult African green monkeys (4 males, and 4 females, body weight 3.5-6kg) were inoculated via a combination of intranasal (0.5 ml per nostril), intratracheal (4 ml), oral (1 ml) and ocular (0.25 ml per eye) of a 4⨯10^5^ TCID50/ml (3⨯10^8^ genome copies/ml) virus dilution in sterile DMEM. Two control animals (1 male and 1 female, body weight 4.5-5.5kg) were inoculated via the same routes with the same dose and volume of inoculum that was gamma-irradiated to render the virus noninfectious (Feldmann et al., 2019). The animals were observed twice daily for clinical signs of disease using a standardized scoring sheet (Munster et al., 2020); the same person assessed the animals throughout the study. The predetermined endpoint for this experiment was 3 days post inoculation (dpi) for the two control animals that were inoculated with irradiated virus and one group of 4 animals inoculated with infectious SARS-CoV-2, and 10 dpi for the remaining 4 animals inoculated with infectious SARS-CoV-2. Clinical exams were performed on 0, 1, 3, 5, 7, and 10 dpi on anaesthetized animals. On exam days, clinical parameters such as bodyweight, body temperature and respiration rate were collected, as well as ventro-dorsal and lateral chest radiographs. Blood and nasal, throat, and rectal swabs were collected during all clinical exams. Additionally, during on 1, 3, and 5 dpi animals were intubated and bronchoalveolar lavages were performed using 10-20ml sterile saline. After euthanasia, necropsies were performed. The percentage of gross lung lesions was scored by a board-certified veterinary pathologist and samples of the following tissues were collected: cervical lymph node, conjunctiva, nasal mucosa, nasal septum, oropharynx, tonsil, trachea, all six lung lobes, mediastinal lymph node, right and left bronchus, heart, liver, spleen, kidney, stomach, duodenum, jejunum, ileum, cecum, colon, and urinary bladder. Histopathological analysis of tissue slides was performed by a board-certified veterinary pathologist blinded to the group assignment of the animals.

### Virus and cells

SARS-CoV-2 isolate nCoV-WA1-2020 (MN985325.1) (Harcourt et al., 2020) (Vero passage 3) was kindly provided by CDC and propagated once in VeroE6 cells in DMEM (Sigma) supplemented with 2% fetal bovine serum (Gibco), 1 mM L-glutamine (Gibco), 50 U/ml penicillin and 50 μg/ml streptomycin (Gibco) (virus isolation medium). The used virus stock was 100% identical to the initial deposited genbank sequence (MN985325.1) and no contaminants were detected. This virus was gamma-irradiated with a dose of 2MRad to produce a non-infectious inoculum (Feldmann et al., 2019). Absence of infectious virus after gamma-irradiation was confirmed in Vero E6 cells. VeroE6 cells were maintained in DMEM supplemented with 10% fetal calf serum, 1 mM L-glutamine, 50 U/ml penicillin and 50 μg/ml streptomycin.

### Quantitative PCR

RNA was extracted from swabs and BAL using the QiaAmp Viral RNA kit (Qiagen) according to the manufacturer’s instructions. Tissues (30 mg) were homogenized in RLT buffer and RNA was extracted using the RNeasy kit (Qiagen) according to the manufacturer’s instructions. For detection of genomic and subgenomic RNA, 5 µl RNA was used in a one-step real-time RT-PCR assay (Corman et al., 2020; Wolfel et al., 2020) using the Rotor-Gene probe kit (Qiagen) according to instructions of the manufacturer. In each run, standard dilutions of counted RNA standards were run in parallel, to calculate copy numbers in the samples.

### Virus titration and isolation

Virus titrations were performed by end-point titration in Vero E6 cells. Cells were inoculated with 10-fold serial dilutions of swab and BAL samples. Virus isolation was performed on tissues by homogenizing the tissue in 1ml DMEM and inoculating Vero E6 cells in a 24 well plate with 250 µl of cleared homogenate and a 1:10 dilution thereof. One hour after inoculation of cells, the inoculum was removed and replaced with 100 µl (virus titration) or 500 µl virus isolation medium. Six days after inoculation, CPE was scored and the TCID50 was calculated.

### Histopathology

Histopathology, immunohistochemistry and in situ hybridization were performed on African green monkey tissues. After fixation for a minimum of 7 days in 10 % neutral-buffered formalin and embedding in paraffin, tissue sections were stained with hematoxylin and eosin (HE).

Immunohistochemistry was performed using a custom-made rabbit antiserum against SARS-CoV-2 N at a 1:1000 dilution, using a CD68 clone KP1 mouse monoclonal antibody (Dako/Agilent cat #M0814) at a 1:100 dilution to identify macrophages, and using Cytokeratin clone AE1/AE3 mouse monoclonal antibody (Dako/Agilent cat #M3515) at a 1:100 dilution to identify epithelial cells. In situ hybridization was used for detection of SARS-CoV-2 RNA in selected whole tissue sections of the lungs using the RNAscope VS Universal AP assay (Advanced Cell Diagnostics Inc.) as described previously (de Wit et al., 2013) and using probe directed against the SARS-CoV-2 S gene (cat# 848569). Stained slides were analyzed by a board-certified veterinary pathologist.

### Single-Cell Sequencing of lung and mediastinal lymph nodes

Lung sections and mediastinal lymph nodes were taken at the time of necropsy of the animals and processed. Cell suspensions were generated by manually dicing tissue, enzymatically digesting in RPMI containing 0.1 mg/ml Liberase TM (Sigma 5401127001) and 0.02 mg/ml DNaseI (Sigma 11284932001) at 37°C, and then passing through a 100 um filter. Suspensions were subjected to ACK lysis and final washes in PBS containing 0.1% MACS BSA (Miltenyi 130-091-386). 10,000 cells were prepared for 10X Genomics gel bead emulsions. The 10X genomics version 3.0 chemistry was used. cDNA for the individual cells was generated and libraries prepped according to the manufactures protocol. After the final libraries were generated, samples were inactivated for any potentially remaining virus using 500ul of AVL with 500ul of ethanol with a sample volume of 140ul. After a minimum 10-minute incubation, samples were removed from the high-containment laboratory following standard protocols and the libraries were extracted from the AVL using the Qiagen AllPrep DNA spin columns (Cat 80204). Samples were then sent for quantification and sequencing. Samples were sequenced on the NextSeq550 using the 10X suggested cycling.

### Processing of single-cell sequencing samples

Samples were process through the cellRanger pipeline to perform de-multiplexing and generate count tables. Alignment was done against the African Green Monkey Ensembl genome (ChlSab1.1) with the SARS-CoV-2 genome (NC_045512.2) included to be able to parse out reads associated with the viral genome. Samples were then read into R (V3.6.2) using Seurat (V3.1.5) (Stuart et al., 2019). Since samples were collected across two different days (day 3 and day 10 post infection), we wanted to account for potential batch effects in the global dataset. For this reason, we then integrated the samples using the IntegrateData function. Cells were filtered that contained abnormally high mitochondrial genes (greater than 3 standard deviations above the median) and cells that are likely doublets were re-labeled (ratio of unique features to UMI per cell < 0.15). Also, cells containing less than or greater than 3 sd of UMI compared to the population total were removed to filter for noise. Finally, the PCA and UMAP projections were calculated for the samples and significant clusters of cells were identified. The lung and the lymph node samples were analyzed separately. Gene set enrichment analysis was performed using fgsea (Sergushichev, 2016) and the MSig DB (Subramanian et al., 2005) c2cp gene sets.

### Cell type identification for single-cell sequencing data

To determine the identity of either clusters of cells or individual cells, we developed an unbiased method that uses a transcriptional profile of cells instead a few known marker genes. For the reference data, we used an annotated single cell sequencing dataset from (Madissoon et al., 2019). For each of the cell type present in the dataset of lung or spleen tissue, we calculated the differential gene expression using the FindMarkers function in Seurat. To find genes strongly associated with each individual cell type we filtered the data to contain only those genes with an average logFG greater than 1 and where the difference in the percentage of cells in the cell type of interest expressing the gene compared to the percentage of cells in all other cell types expressing the gene was greater than 0.5. We used this gene set in either the lung or spleen of the human samples to develop the marker gene set and calculated the average expression of the marker gene set in each cell type. This generated a matrix of the marker genes to cell types. Then a correlation of the marker genes in the annotated data was compared to the individual cell or cluster in the African green monkey data. This generated a score for the unknown cell or cluster to a known annotation. Using this method, we found that most clusters contained predominately just 1 cell type (Fig. 4E,F). This is a similar method that was developed for the mouse cell atlas (Han et al., 2018). The results were validated by looking at the expression of the marker genes across the different cell annotations (Fig. 3B and Fig. 5B). When determining the identity on an individual cell level, an additional step was added to help correct falsely identified cells. Using the knn graph generated in the FindNeighbors function in Seurat, for each cell, its closest neighbors were determined. Once the nearest neighbors were determined, the identities of these neighbors were pulled out. If > 70% of the nearest neighbors had one specific identity, the cell identity was reassigned as such. This was run multiple times until a stable number of unidentified cells was found (determined by small changes in unknown cell identities). For those that were not able to be identified, the identity from the original transcriptional profile was used.

Finally, to identify clusters that are specifically cells undergoing rapid cell division, we used the CellCycleScore function in Seurat to identify which cell cycle each cell was likely in. We then determined that clusters where greater than 95% of the cells were in G2M or S phase were dividing clusters and were labeled as such.

### SARS-CoV-2 read enrichment

To analyze the enrichment of reads across the SARS-CoV-2 genome, we used Integrative Genome Viewer (Robinson et al., 2011) to find read pile-ups. Cells were labeled as positive for viral RNA if they contained any counts to the viral genome.

### Clustering biases in scRNA-Seq Data

To determine if there is a clustering bias between two cell types, a new method was developed. Across any cluster of cells, the dataset is subset and re-normalized internally to that cluster. Then the significant principal components (PC) (containing up to 99% of the variance explained) are calculated. Along each principal component, the location of the cells is pulled and grouped based on the two conditions that are being compared. The median location of each of the cell populations along the PC is calculated and the distance measured. This is carried out for all PCs and all clusters. To identify outliers with the strongest clustering bias, points outside the mean and 2 standard deviations across all the PCs and cell types are noted.

### Statistical Analysis

Statistical tests comparing cell numbers were carried out in Prism V8 using an ANOVA. Statistical tests for gene expression in single cell data was carried in Seurat V3.6. Graphs were generated either in ggplot2 (Wickham, 2016) or Prism V8.

### Data availability

Animal model data have been deposited in Figshare: https://doi.org/10.6084/m9.figshare.12818726.v1. RNA sequencing data have been deposited in NCBI’s Gene Expression (Edgar et al., 2002) and are accessible through GEO Series accession number GSExxx (https://www.ncbi.nlm.nih.gov/geo/query/acc.cgi?acc=GSExxx).

## Notes

https://doi.org/10.6084/m9.figshare.12818726.v1

### Competing Interest Statement

The authors have declared no competing interest.

