## Supplemental Data for "SARS-CoV-2 infection dynamics in lungs of African green monkeys"

*[1]Laboratory of Virology, [2]Rocky Mountain Veterinary Branch and [3]Research Technologies Branch, Division of Intramural Research, National Institute of Allergy and Infectious Diseases, National Institutes of Health, Hamilton, MT, United States of America, [4]Laboratory of Immune System Biology, Lymphocyte Biology Section, Division of Intramural Research, National Institute of Allergy and Infectious Diseases, National Institutes of Health, Bethesda, MD, United States of America*

**Table S1.** Virus isolation from tissues of African green monkeys inoculated with SARS-CoV-2 and euthanized at 3 and 10 dpi.

| | AGM3 | AGM4 | AGM5 | AGM6 | AGM7 | AGM8 | AGM9 | AGM10 |
|---|---|---|---|---|---|---|---|---|
| | 3 dpi | | | | 10 dpi | | | |
| RUL | + | ++ | - | + | - | - | - | - |
| RML | - | - | - | ++ | - | - | - | - |
| RLL | ++ | ++ | ++ | ++ | - | - | - | - |
| LUL | + | - | - | ++ | - | - | - | - |
| LML | - | - | + | ++ | - | - | - | - |
| LLL | ++ | ++ | ++ | ++ | - | - | - | - |
| Duodenum | ND | ND | ND | ND | ND | - | ND | ND |
| Jejunum | ND | ND | ND | ND | ND | - | ND | ND |
| Ileum | ND | ND | ND | ND | ND | ++ | ND | ND |
| Cecum | ND | ND | ND | ND | ND | ++ | ND | ND |
| Colon | ND | ND | ND | ND | ND | - | ND | ND |

+ positive in virus isolation; ++ positive in virus isolation at 1:10 dilution of tissue; - negative in virus isolation; ND not done; RUL right upper lung lobe; RML right middle lung lobe; RLL right lower lung lobe; LUL left upper lung lobe; LML left middle lung lobe; LLL left lower lung lobe

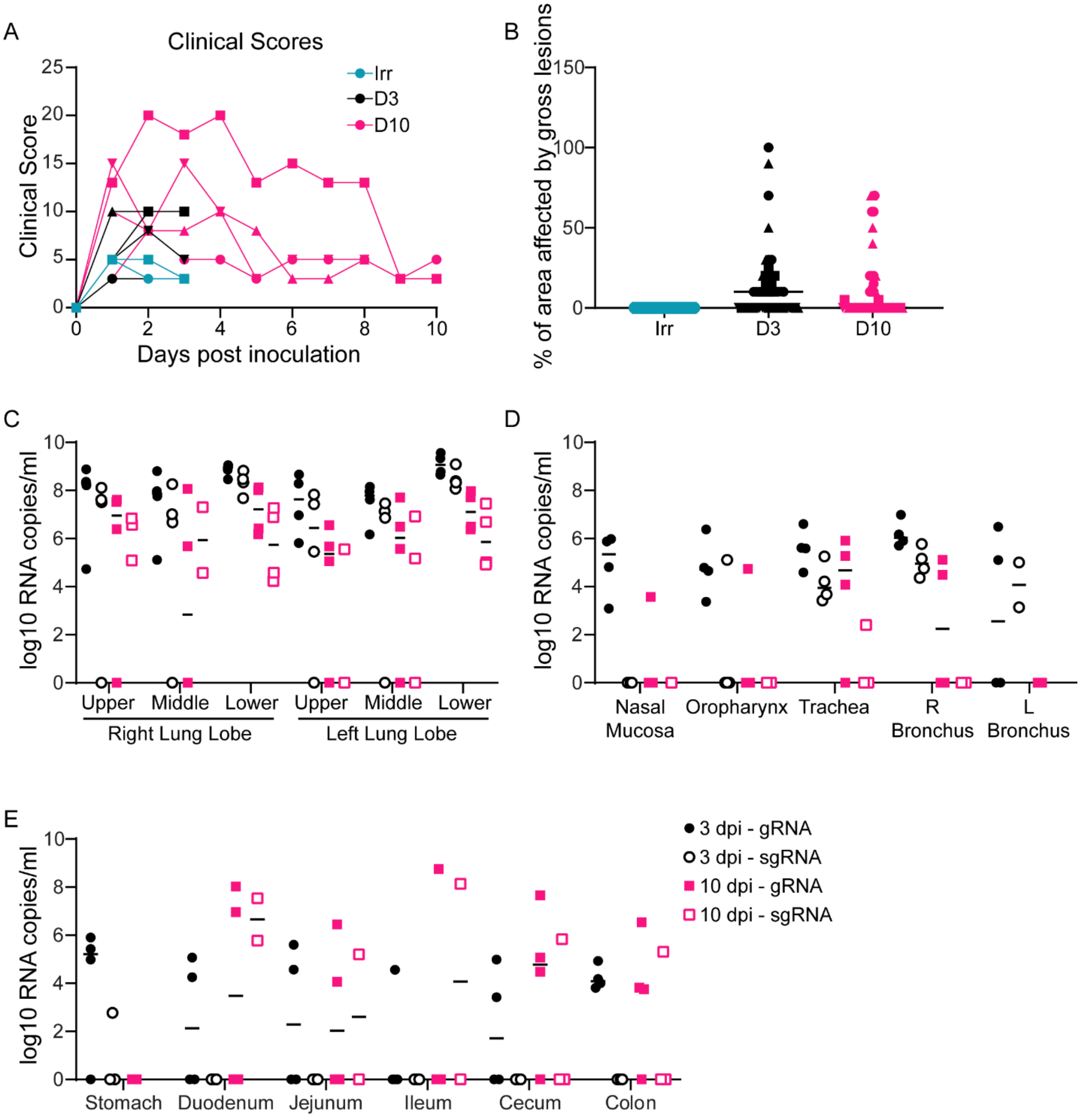


**Figure S1. Clinical scores, gross lung lesions and viral loads in respiratory and gastrointestinal tract of African green monkeys inoculated with SARS-CoV-2.** Two African green monkeys were inoculated with gamma-irradiated SARS-CoV-2 and euthanized at 3 dpi; eight animals were inoculated with infectious SARS-CoV-2 isolate nCoV-WA1-2020 and four of those were euthanized at 3 dpi, and four at 10 dpi. A. After inoculation, animals were observed for disease signs and scored according to a pre-established clinical scoring sheet. B. At necropsy, the percentage of the area of the lungs affected by gross lesions was estimated by a board-certified veterinary pathologist. C. Tissue samples were collected at necropsy and analyzed for the presence of genomic (closed symbols) and subgenomic (open symbols) RNA by qRT-PCR. Teal: animals inoculated with gamma-irradiated virus; black: animals inoculated with infectious virus and euthanized at 3 dpi; pink: animals inoculated with infectious virus and euthanized at 10 dpi. Where indicated, identical symbols have been used to identify individual animals throughout the figures in this manuscript.

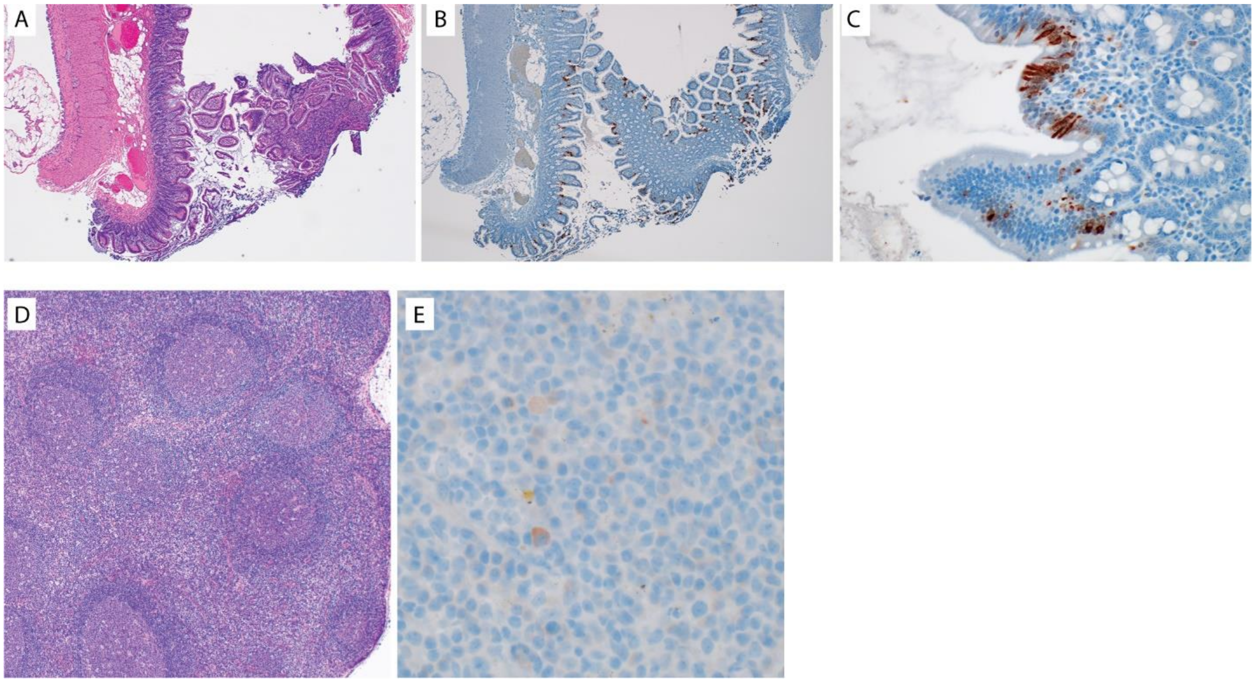


**Figure S2. Histological changes in intestinal tract and lymp nodes of African green monkeys infected with SARS-CoV-2.** A-C. Evidence of virus replication in the ileum of AGM8. A. Histologically, the ileum appears normal. B Viral antigen is detected in cells distributed throughout the mucosa and lamina propria. C. Higher magnification showing viral antigen in the cytoplasm of multiple epithelial cells and in scattered mononuclear cells within the lamina propria. D-E. Evidence of lymph node hyperplasia and virus antigen at 10 dpi. D. Histologically, the lymph node appears to have mild to moderate hyperplasia. Follicles have large germinal centers surrounded by a thin mantel and extend into the center of the node. E. Only rare mononuclear cells stained positive for viral antigen in the cytoplasm. Magnification A, B, D 40x, C, E 400x

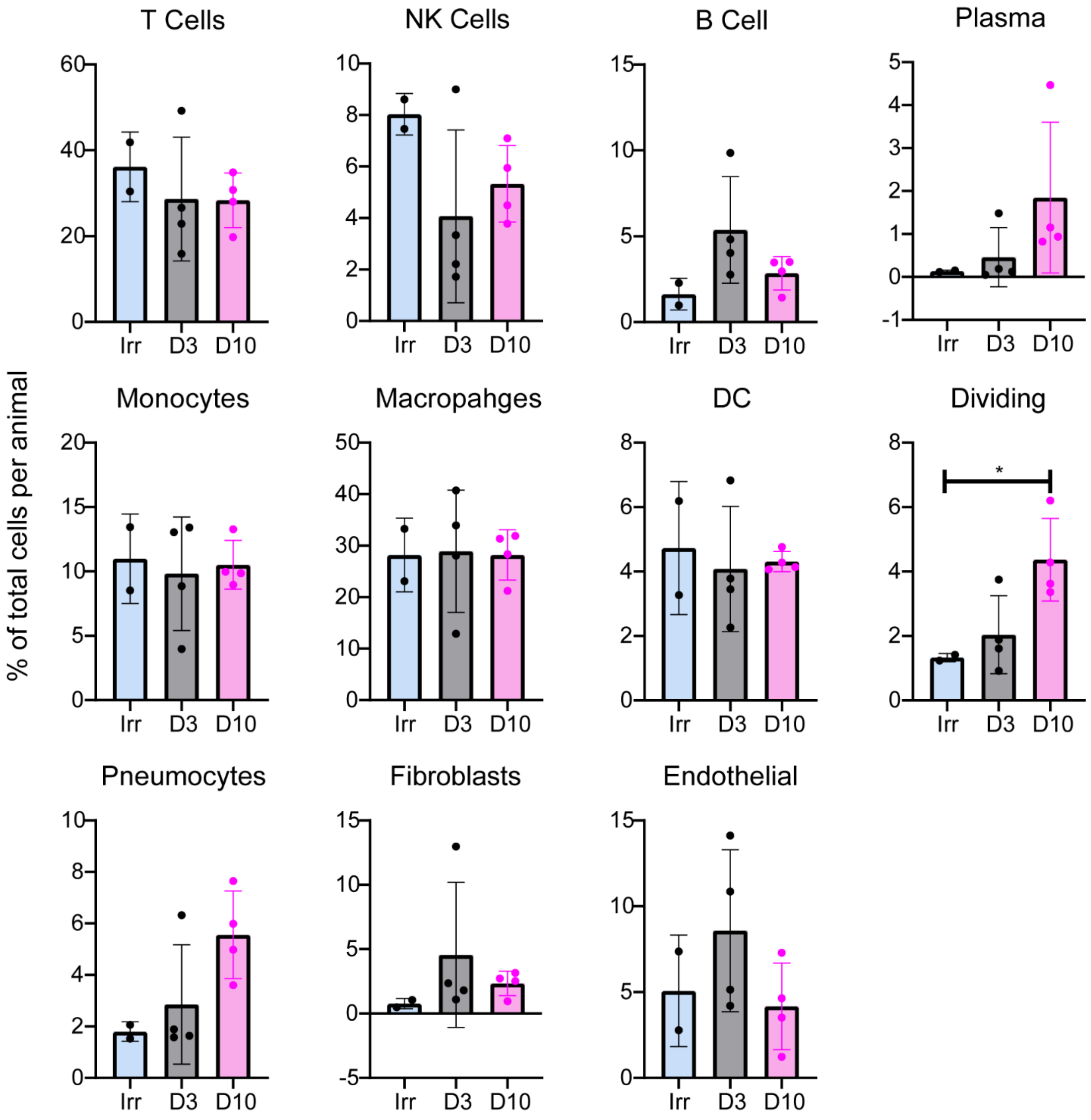


**Figure S3: Percentage of cell types in the lung determined by scRNA-sequencing.** Each graph represents a specific cell annotation. Irr: animals inoculated with gamma-irradiated virus and euthanized at 3 dpi: D3 animals inoculated with SARS-CoV-2 and euthanized at 3 dpi; D10: animals inoculated with SARS-CoV-2 and euthanized at 10 dpi. Each point represents an individual animal. Bars represent the mean of the groups and the error bars indicate standard deviation. Dividing cells were significantly different between the animals inoculated with gamma-irradiated virus and animals inoculated with SARS-CoV-2 and euthanized at 10 dpi ($p = 0.044$) as determined by an ANOVA.

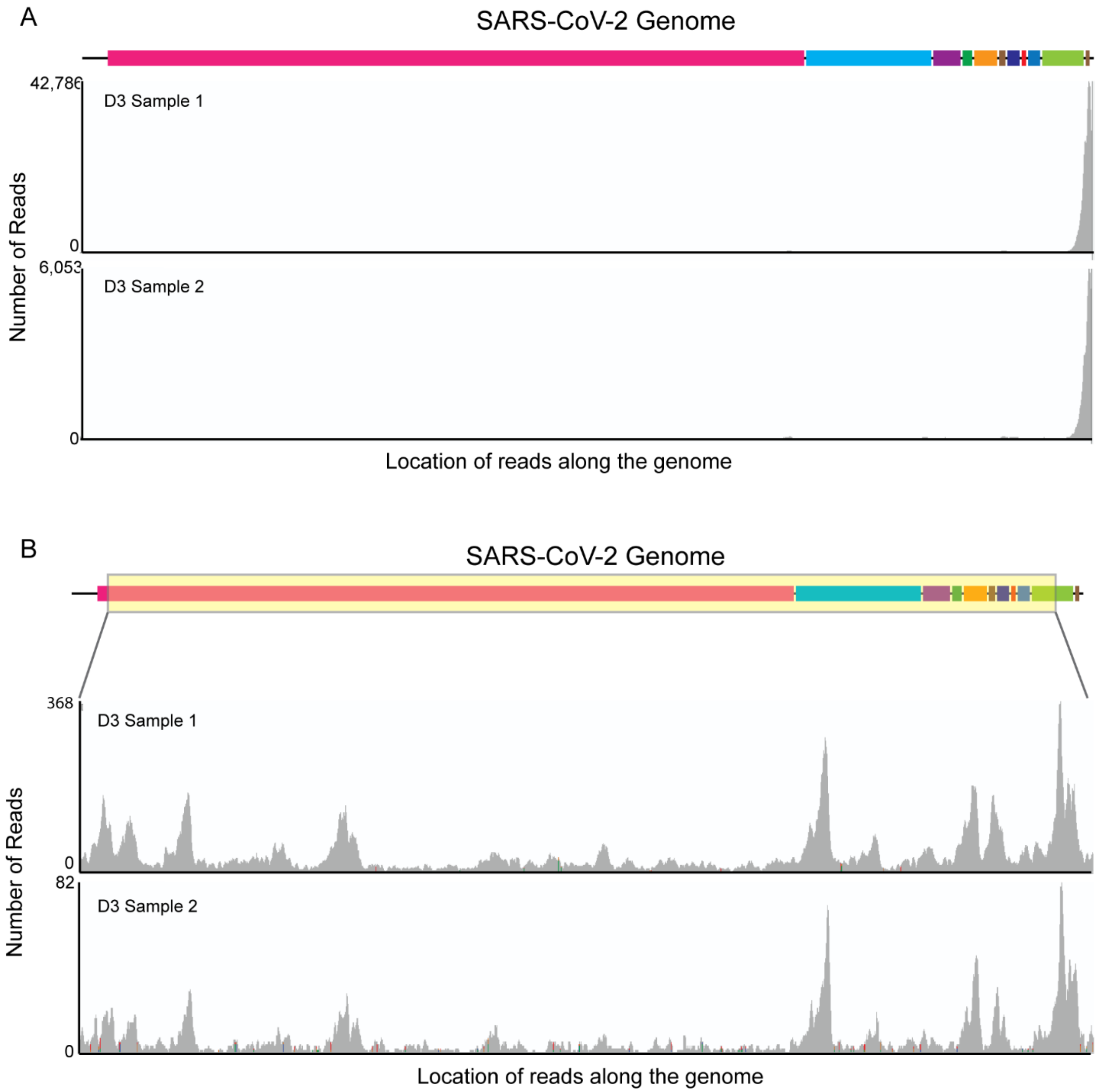


**Figure S4: Coverage of reads from the pseudo-bulk data across the SARS-CoV-2 genome.** A. Histogram of read alignment locations across the SARS-CoV-2 full genome. Two samples from at 3 dpi are represented as they had the most reads to the virus. The x-axis is the location along the genome with the left being the 5′ end and the right being the 3′ end. The y-axis is the number of reads that align at each of the bases. B. A similar graph but for a smaller region of the genome as noted by the yellow highlighted area that excludes the 5′ leader sequence and the 3′ end of the genome.

A

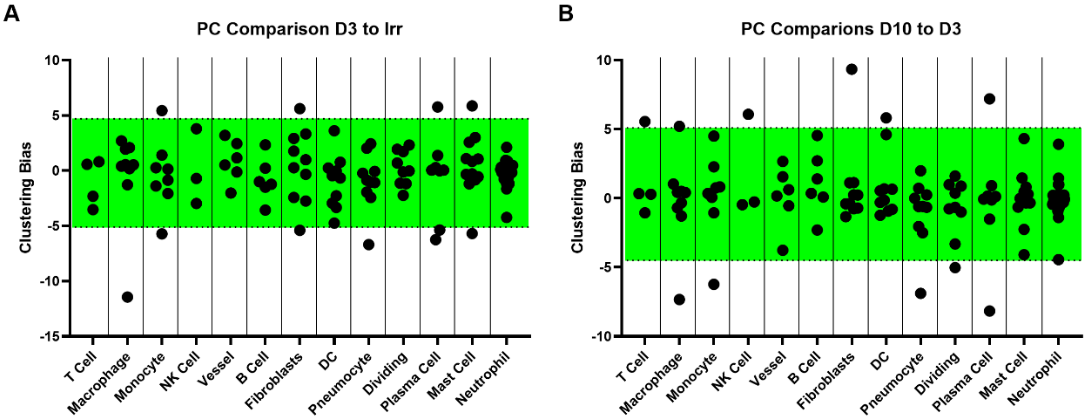


B

C

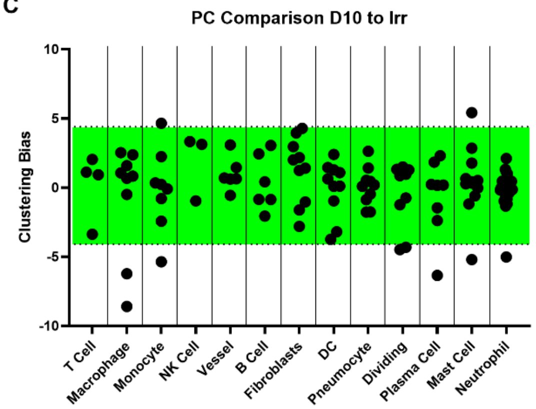


D

| Day 3 Only | Day 3 and Day 10 | Day 10 Only | Day 3 to Day 10 |
|---|---|---|---|
| Monocyte PC1 | Macrophage PC1 | Macrophage PC2 | Macrophage PC2 |
| Fibroblasts PC2 | Monocyte PC3 | Plasma Cell PC5 | Monocyte PC1 |
| Plasma Cell PC2 | | | NK Cell PC1 |
| Pneumocyte PC1 | | | Fibroblasts PC1 |
| | | | Plasma Cell PC1 |
| | | | Plasma Cell PC2 |
| | | | Pneumocyte PC1 |

**Figure S5. Clustering bias across cell populations in the lungs.** A. Clustering bias analysis of animals inoculated with gamma-irradiated virus and euthanized at 3 dpi compared to animals inoculated with SARS-CoV-2 and euthanized at 3 dpi. The x-axis shows the various cell populations identified in the lung. The y-axis shows the clustering bias determined by the difference between the median values of the locations along the principal component. Each point is a significant principal component for a given cell type. The green box represents the data that is within expected noise. Points outside the green box are greater than 2 standard deviations from the mean value. B. The same analysis, now comparing the animals inoculated with SARS-CoV-2 and euthanized at 3 and 10 dpi. C. The same analysis comparing animals inoculated with gamma-irradiated virus and euthanized at 3 dpi to animals inoculated with SARS-CoV-2 and euthanized at 10 dpi. D. A summary table of the analysis. Each column represents if the cell type and PC combination was enriched in SARS-CoV-2 inoculated animals at 3 dpi only (Day 3 only), was enriched in SARS-CoV-2-infected animals compared to animals inoculated with gamma-irradiated virus (Day 3 and Day 10), enriched in SARS-CoV-2-infected animals at 10 dpi only (day 10 only) or enriched when comparing SARS-CoV-2-infected animals at 3 and 10 dpi (Day 3 to Day 10).

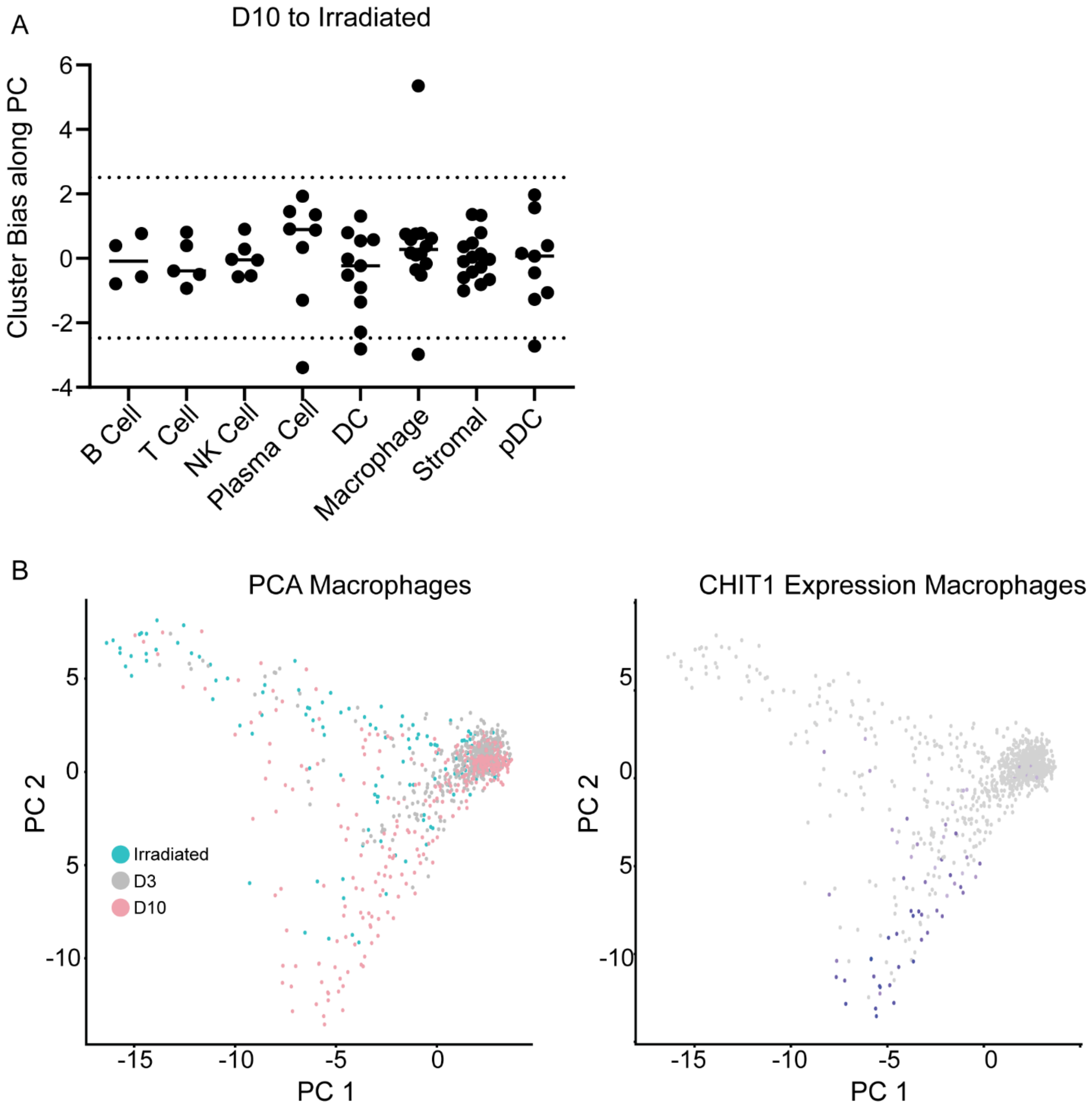


**Figure S6: Ln Comparison of different cell populations.** A. Unbiased cluster bias values (y-axis) of the different cell populations (x-axis) along the different principal components comparing day 10 post inoculations with SARS-CoV-2 to day 3 post inoculations with the irradiated virus. B. PCA projection for the macrophage populations comparing PC1 (x-axis) and PC2 (y-axis). Each point is an individual cell. Left graph is colored based on the sample type (blue = irradiated, grey = day 3 post inoculation, pink = day 10 post inoculation). The right is the same graph but colored based on CHIT1 expression. The darker the purple, the higher the expression value.
